# Perceived time drives physical fatigue

**DOI:** 10.1101/2025.07.13.664573

**Authors:** Pierre-Marie Matta, Julien Duclay, Robin Baurès, Andrea Alamia

**Affiliations:** Univ Toulouse, CNRS, CerCo, Toulouse, France; Univ Toulouse, Inserm, ToNIC, Toulouse, France

**Keywords:** Time deception, False-clock paradigm, Frontal oscillations, Fatigue, Mind-body connection

## Abstract

Recent studies suggest that fundamental physiological processes, such as physical fatigue, rely on perceived rather than actual time. However, the neural correlates underlying this effect and its disentanglement from motivational factors (i.e., performance goals) remain unknown. To investigate the time deception effect on fatigue, we developed a novel EEG design in which participants (N = 24) performed 100 isometric contractions at a fixed pace and resistance in four distinct sessions. The actual contraction duration (short or long) and the calibration of the displayed clock (normal or biased toward acceleration or deceleration) were independently manipulated across sessions to examine whether fatigue and its neural correlates evolved in response to perceived or actual time. Our results show an accumulation of physical fatigue that follows the perceived time, irrespective of motivational factors. This effect was consistently observed only when the clock was slowed down. This time-deception effect involved frontal theta- and beta-band dynamics: theta modulated only under the slowed clock and beta robustly shaped by perceived time across both slowed and accelerated clocks, while motor beta showed no modulation. Further analyses highlighted the key role of frontal oscillatory dynamics in the effectiveness of the time-deception effect on physical fatigue.

**Significance statement:** Can a clock change the course of physical fatigue? In this study, we addressed this question while controlling for motivational confounds and monitoring EEG-associated power modulations. Our findings demonstrate a slowed-down fatigue accumulation in the presence of a slowed-down clock. This time manipulation effect was driven by a frontal oscillatory dynamic that largely followed the perceived time. These results highlight the direct influence of psychological factors on physiological processes and unveil the neural correlates underlying this effect.

## Introduction

The relationship between the mind and body has been a subject of inquiry for centuries (Dijkerman & Lenggenhager, 2018), raising fundamental questions about how mental states influence bodily processes. A prevalent approach to exploring this relationship involves deceptively manipulating participants’ beliefs while recording psychophysiological variables (Crum et al., 2011; Crum & Langer, 2007; H. S. Jones et al., 2013; Williams et al., 2014). Among the various psychological parameters that can be successfully deceived, manipulating the subjective passage of time has been suggested as particularly relevant because of the trust we routinely place in its measurement (Thönes et al., 2018). This manipulation is typically achieved by covertly altering the clock’s speed displayed to participants during a task, either slowing it down or accelerating it (Thönes et al., 2018). For instance, when the passage of time is artificially slowed down, studies have reported reduced perceived pain in response to thermal stimulation (Pomares et al., 2011) but also a smaller decrease in blood glucose level (Park et al., 2016) and slower healing (Aungle & Langer, 2023). Similarly, studies interested in physical activity have shown that, in the presence of a slowed-down clock, there is a slower spontaneous motor activity (Craik & Sarbin, 1963) and a slower accumulation of physical fatigue (Matta et al., 2024). Conversely, accelerating the perceived passage of time has been associated with opposite effects, with the notable exception of physical fatigue, which, to our knowledge, remains unexplored in this context.

Physical fatigue, broadly defined as the decline in the ability to generate force or sustain performance (Bigland-Ritchie & Woods, 1984; Gandevia, 2001), is a central determinant of endurance, productivity, and daily functioning (Enoka & Duchateau, 2016). Beyond its role in limiting athletic and occupational performance, it is also closely linked to injury risk (Chan, 2011; Cunningham et al., 2022; Ibrahim et al., 2023; C. M. Jones et al., 2017; Kreher & Schwartz, 2012; Rodrigues et al., 2023) and impaired quality of life (Behrens et al., 2023). Understanding how it may be shaped by psychological factors is therefore of both theoretical and practical significance, with implications for athletes, workers in physically demanding professions, and individuals in everyday life. This makes physical fatigue a particularly relevant phenomenon for investigating deceptive manipulations. Furthermore, physical fatigue develops progressively as a function of task duration, reflecting the cumulative physiological and neural demands imposed by sustained activity (Bigland-Ritchie & Woods, 1984; Edwards, 1981; Enoka & Duchateau, 2008; Gandevia, 2001). As a result, fatigue is intrinsically linked to temporal representations, as information about elapsed and remaining time provides a key reference for regulating effort throughout a task. In most goal-directed behaviors, time constitutes a fundamental performance constraint, shaping pacing strategies, effort allocation, and expectations about task completion (Brehm & Self, 1989; de Koning et al., 2011; Noakes, 2011; Tucker, 2009). Consequently, beliefs about time serve as a critical reference signal during sustained physical activity, pointing out the importance of examining how these temporal representations may influence the development of physical fatigue.

Nevertheless, important gaps remain in our understanding of how time perception influences physical fatigue: although these perceived time-dependent effects have been demonstrated for several variables (Thönes et al., 2018), their underlying neural mechanisms remain poorly understood. Moreover, motivational factors, such as performance goals, are critical confounds often unaccounted for when measuring deceptive effects. These confounds are particularly important in physical fatigue, in which previous studies have consistently employed deceptive manipulations to enhance physical performance (i.e., the amount of time a task-to-exhaustion is maintained or the time needed to complete a given distance) (Ansdell et al., 2018; Ducrocq et al., 2017; Matta et al., 2024). Consequently, previous results on physical fatigue may be alternatively explained by performance-related goals rather than time perception.

In the current study, we investigate the neural correlates of the time deception effect on the accumulation of physical fatigue and carefully isolate this effect from motivational confounds. In addition, we also test its temporal symmetry, that is, when the clock is either slowed down or accelerated. To this end, we used an original 2×2 design, in which each session consisted of 100 isometric contractions at a given pace against a fixed, individualized resistance. The contraction’s actual duration (short or long) and the calibration of the displayed clock (normal or biased toward acceleration or deceleration) were manipulated between sessions to test whether physical fatigue was evolving according to the perceived (i.e., displayed) or actual time (Fig. 1A). Unlike previous studies, we tested all four possible combinations for the first time in a within-subject design. We recorded EMG and EEG signals to assess fatigue accumulation and associated neural correlates. We specifically focused on oscillatory dynamics at the sensor level in the motor and prefrontal cortex, as both regions are involved in physical fatigue and effort maintenance (Bigliassi & Filho, 2022; Matta et al., 2025; McMorris et al., 2018; Robertson & Marino, 2016).

**Figure 1.**
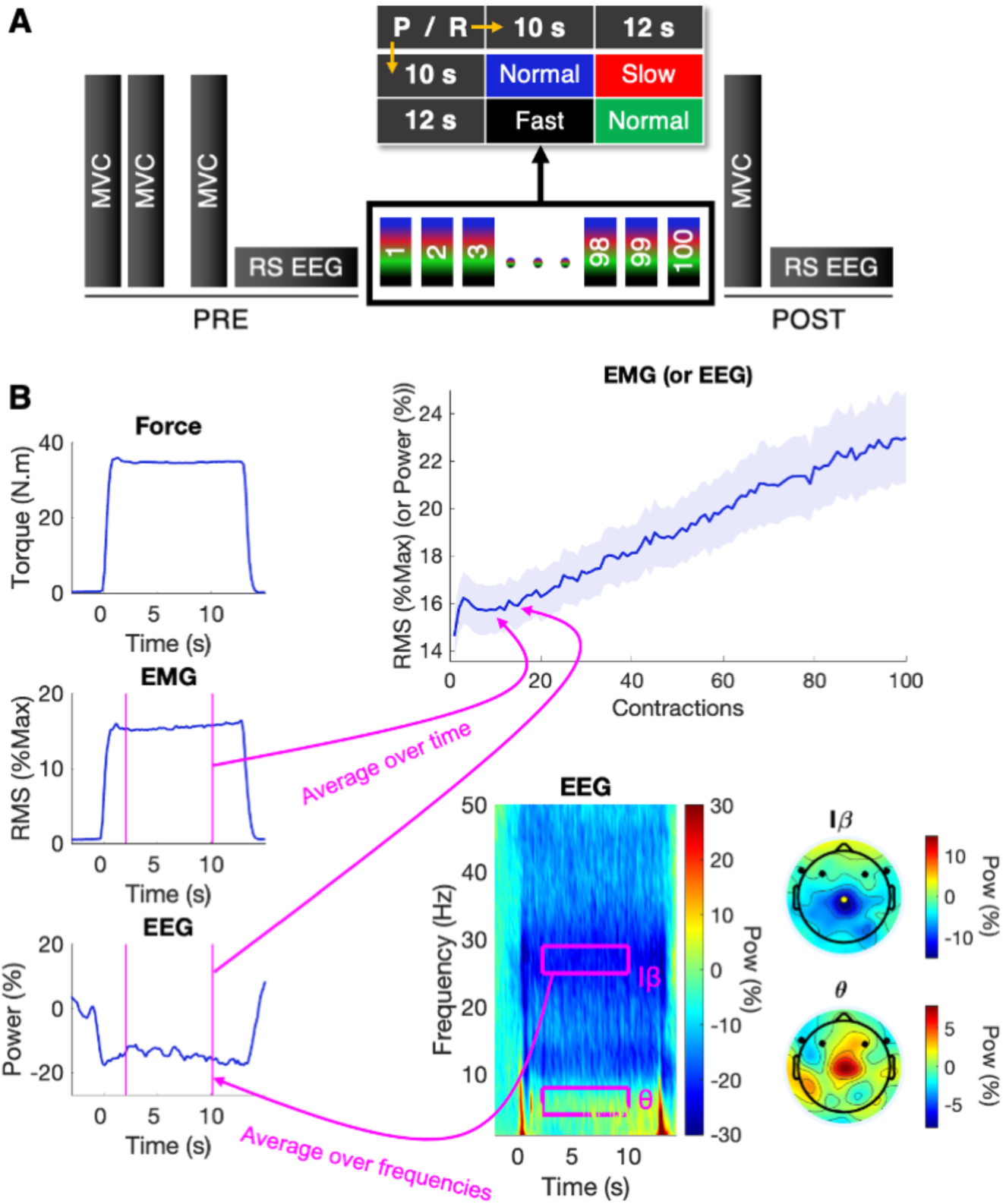
Experimental protocol and EMG/EEG computation. A. Experimental Protocol. All participants performed 100 contractions during four separate sessions. While the rest time between contractions was identical (5s) for each session, each contraction’s real (R) and perceived (P) time were manipulated. In each session, the contractions’ time was either short (10s) or long (12s), and the digital clock displayed in front of the participants was manipulated to induce either a short (10 s) or long (12 s) perceived duration. This resulted in a within-subject, partially crossed four-condition design manipulating Real duration (10 s vs. 12 s) and Perceived duration (10 s vs. 12 s), yielding four counterbalanced conditions: P10R10 (normal clock), P12R12 (normal clock), P10R12 (slowed clock), and P12R10 (fast clock). Before and after the task, Maximal Voluntary Contractions (MVC) were performed to assess fatigue induced by the 100 contractions. Resting State (RS) EEG (3 min) was recorded before and after the task to quantify the associated EEG correlates. B. EMG/EEG Computation. All panels represent the typical trace of a single contraction, except the upper-right panel, which illustrates the entire task. EMG and EEG were recorded throughout the task and explicitly analyzed over a 2-10 s window centered on each contraction (0 s = contraction onset), from which we extracted a single average value per contraction. The EMG was computed in normalized Root Mean Square (RMS), while we computed the mean power of the theta (θ) and an individualized beta-band (Iβ) for the EEG, in frontal electrodes and Cz (illustrated in black and yellow respectively on the topographical plots) (see Methods for details).

## Results

### Design Summary

Participants performed four sessions (each spaced 3 to 10 days apart) of 100 quadriceps isometric contractions, during which EMG and EEG were continuously recorded. For each participant, the contraction intensity was identical across sessions (≤ 20% of the participant’s maximal force; see Methods for details). While the rest time between contractions was identical for each session, each contraction’s real (R) and perceived (P) time were independently manipulated. Each contraction could be either short (10 s) or long (12 s) in real time, and the digital clock displayed in front of the participants was manipulated to induce either a short (10 s) or long (12 s) perceived duration. This resulted in a 2 × 2 design, with the factors Real duration (10 s vs. 12 s) and Perceived duration (10 s vs. 12 s), yielding four counterbalanced conditions: P10R10 (normal clock), P12R12 (normal clock), P10R12 (slowed clock), and P12R10 (fast clock) (Fig. 1). Before and after the task, participants performed Maximal Voluntary Contractions (MVC) and 3-minute EEG resting states.

### Neuromuscular fatigue

We first assessed the effects of real and perceived time on physical fatigue, quantified as the pre-post change in maximal voluntary torque (MVT). Analysis of the pre-post difference in MVT revealed no significant main effect of perceived time (p = .90), real time (p = .47), nor their interaction (p = .23). This lack of effect may stem from the relatively small variation in workload across sessions (Matta et al., 2025) combined to a limited sensitivity of pre-post MVT measurements to accurately detect physical fatigue (Lebesque et al., 2022).

We next examined whether perceived time modulated fatigue accumulation during the task, quantified as the EMG Root Mean Square (RMS) activity recorded across contractions (Kim et al., 2007; Merletti et al., 1991). Model comparison showed that allowing for a nonlinear effect of contraction number did not improve model fit relative to a linear model, (χ²_(4)_ = 7.10, p = .13), and the linear model was therefore retained. This model revealed a significant effect of contraction number (F_(1, 23)_ = 45.97, p < .0001), as well as significant main effects of perceived time (F_(1, 9546)_ = 16.26, p < .0001). The interaction between contraction number and perceived time was significant (F_(1, 9546)_ = 15.12, p = .0001), as was the interaction between contraction number and real time (F_(1, 9546)_ = 32.08, p < .0001). The three-way interaction between contraction number, perceived time, and real time was also significant (F_(1, 9546)_ = 8.77, p = .003), while no significant main effect of real time alone nor interaction between perceived and real time was observed (all p > .59). Post-hoc contrasts on the estimated marginal slopes indicated that perceived time did not affect EMG rate when real time was short (p = 1, d_z_ = 0.13, BF = 0.22). When real time was long, however, EMG increased significantly less for shorter perceived time (p < .0001), highlighting a strong (d_z_ = 0.99) slowing of fatigue when the clock was slowed. Consistent with this effect, real time did not show a significant effect when perceived time was short (p = .22, d_z_ = 0.39, BF = 0.30). Conversely, when perceived time was long, EMG increased less for short real time (p < .0001), indicating here a strong (d_z_ = 1.24) dependency on the real time when the clock was accelerated (Fig. 2A, 2B). Together, these results suggest that perceived time modulates the rate of physical fatigue accumulation, effectively slowing it when the task feels shorter (Matta et al., 2024), whereas, conversely, fatigue accumulation cannot be accelerated when the task appears longer.

**Figure 2.**
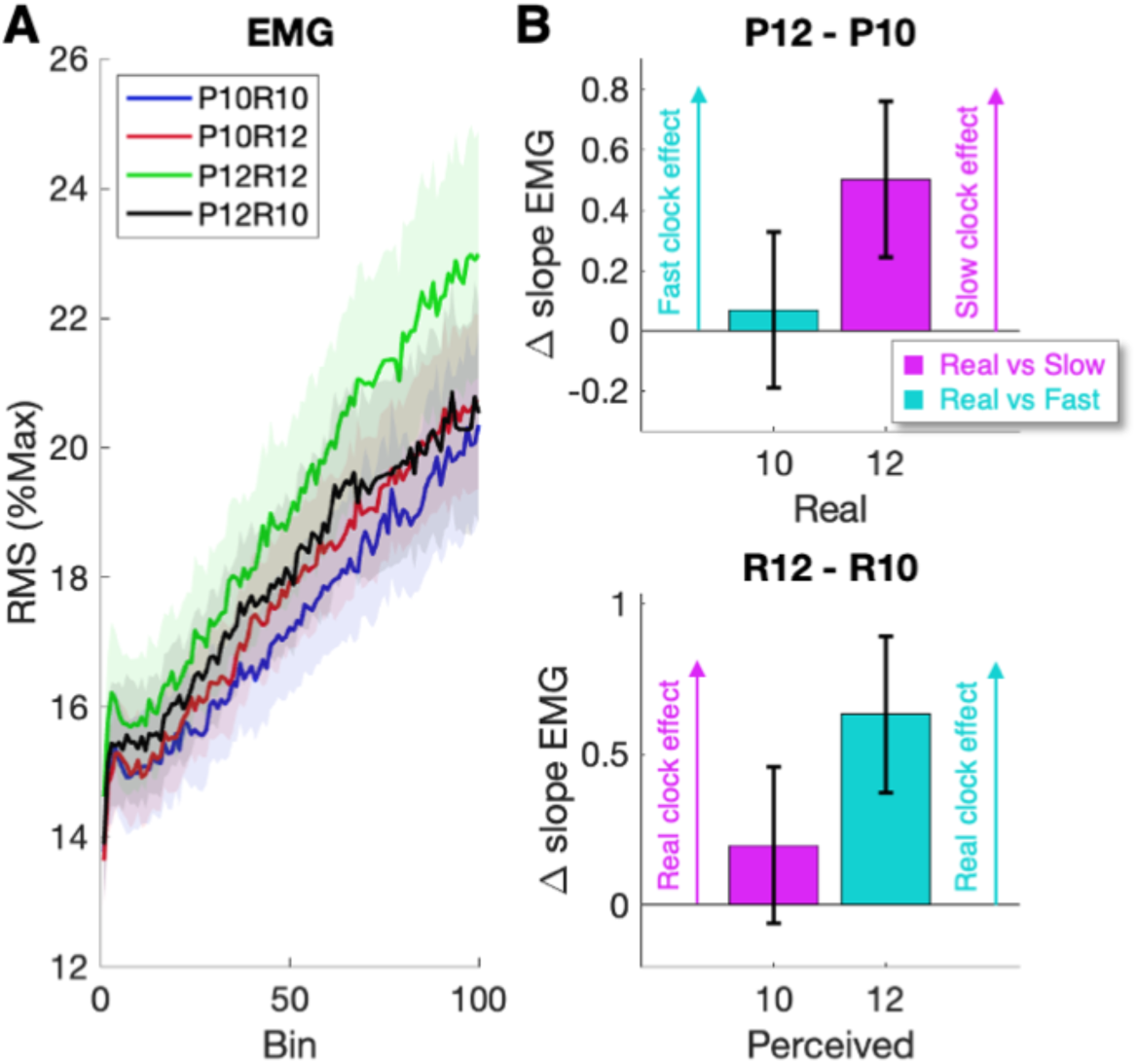
Physical Fatigue. A. EMG modulations over the main task. EMG is expressed in normalized Root Mean Square (RMS) over contractions, whose increase reflects neuromuscular fatigue accumulation. The lines represent the averaged distribution, and the shaded areas are the standard error of the mean (SEM). B. Fatigue differences computed for each contrast. The upper panel illustrates perceived-time effects by holding real duration constant and contrasting long versus short perceived durations, whereas the lower panel illustrates real-time effects by holding perceived duration constant and contrasting long versus short real durations. Both panels represent the difference in slope of the EMG-RMS over the bins. The bar plots represent model-based estimates of condition differences, with error bars indicating 95% confidence intervals. Intervals not crossing zero indicate statistically significant effects at a Bonferroni-corrected alpha of 0.05.

### Power modulations in motor areas

After showing the influence of perceived time on physical fatigue, we investigated the neural correlates underlying this effect. We first examined oscillatory activity in the Cz electrode, commonly used as an approximation of activity in the motor cortex (Dal Maso et al., 2012; Forrester et al., 2006; Gordon et al., 2023; Meier et al., 2008), a key region involved in executing voluntary contractions (Brown, 2000; Pearson, 2000). Focusing on this sensor, we assessed relative spectral power changes in each participant’s individualized beta-band (Iβ) (Matta et al., 2025) (i.e., a 4 Hz spectral window in the beta band with the largest desynchronization, see Methods for details).

We compared two linear mixed-effects models differing in the complexity of the contraction term (natural spline with 1 vs. 2 degrees of freedom). Model comparison indicated that the second-degree spline model provided a better fit (χ²_(4)_ = 11.18, p = .025), and was therefore retained. Analysis of variance on the selected model revealed a significant main effect of contraction number (F_(2, 45.4)_ = 8.01, p = .001), as well as a significant interaction between contraction number and perceived time (F_(2, 9542)_ = 5.28, p = .005), showing a larger decrease in Iβ Cz power when perceived time was shorter. To better understand this effect, we compared the slope differences between the two normal sessions (P10R10 vs. P12R12) but found no difference (p = 0.60, uncorrected, d_z_ = 0.11, BF = 0.23), which may reflect the relatively small discrepancy in workload between the two sessions (Matta et al., 2025) and limits the interpretability of these results. No significant main effects of perceived or real time alone, nor significant interactions involving real time or the three-way interaction, were observed (all p > .05).

We next investigated the effects of perceived and real time on the pre-post difference in Iβ Cz, reflecting overall changes in beta activity across the session. A linear mixed-effects model including participant as a random intercept revealed no significant main effects of perceived time (p = .91) or real time (p = .89), and no perceived × real time interaction (p = .26). This lack of effect may relate to the relatively small discrepancy in workload between the sessions (Matta et al., 2025).

### Power modulations in frontal electrodes

Next, we investigated oscillatory activity at frontal electrodes (F3, F4, F7, F8) – commonly used as an approximation of prefrontal cortex (PFC) activity (Okamoto et al., 2004; Robertson & Marino, 2015), as this region has been proposed to play a key role in effort maintenance (Bigliassi & Filho, 2022; Denis, 2020; McMorris et al., 2018; Robertson & Marino, 2016). We analyzed relative spectral power changes in the theta (θ) and Iβ bands. While frontal θ is assumed to reflect cognitive control (Cavanagh & Frank, 2014), the β band has also been proposed to have a similar function (Stoll et al., 2016).

To assess how perceived time modulates θ frontal activity, we first compared two linear mixed-effects models differing in the complexity of the contraction term (natural spline with 1 vs. 2 degrees of freedom). The more complex model did not improve fit (χ²_(4)_ = 4.91, p = .30), and the linear model was retained. The selected model revealed no significant main effects of perceived time (F_(1, 9546)_ = 0.90, p = .344), real time (F_(1, 9546)_ = 0.59, p = .443), or contraction number (F_(1, 23)_ = 0.39, p = .540). A significant interaction was observed between contraction number and perceived time (F_(1, 9546)_ = 29.65, p < .0001), as well as between perceived and real time (F_(1, 9546)_ = 12.56, p < .0001). The three-way interaction among contraction number, perceived time, and real time was also significant (F_(1, 9546)_ = 5.49, p = .019), while the interaction between contraction number and real time was not significant (p = .65). At short real time, post-hoc contrast on the estimated marginal slopes was inconclusive, showing no reliable difference in the rate of increase in θ frontal for long versus short perceived time (p = .11, d_z_ = 0.45, BF = 0.41). However, when real time was long, the rate of increase in θ frontal was significantly (p < .0001) steeper for long perceived time, with a strong effect size (d_z_ = 1.12). In parallel, real-time influence on θ slopes was inconclusive at short perceived duration (p = .19, d_z_ = 0.40, BF = 0.46) and showed evidence for an absence of difference at long perceived time duration (p = .63, d_z_ = 0.27, BF = 0.28) (Fig. 3A, 3B). Overall, these results indicate that frontal theta changes across the task robustly follow perceived time when the clock is slowed down, while the results are inconclusive regarding the accelerated clock.

**Figure 3.**
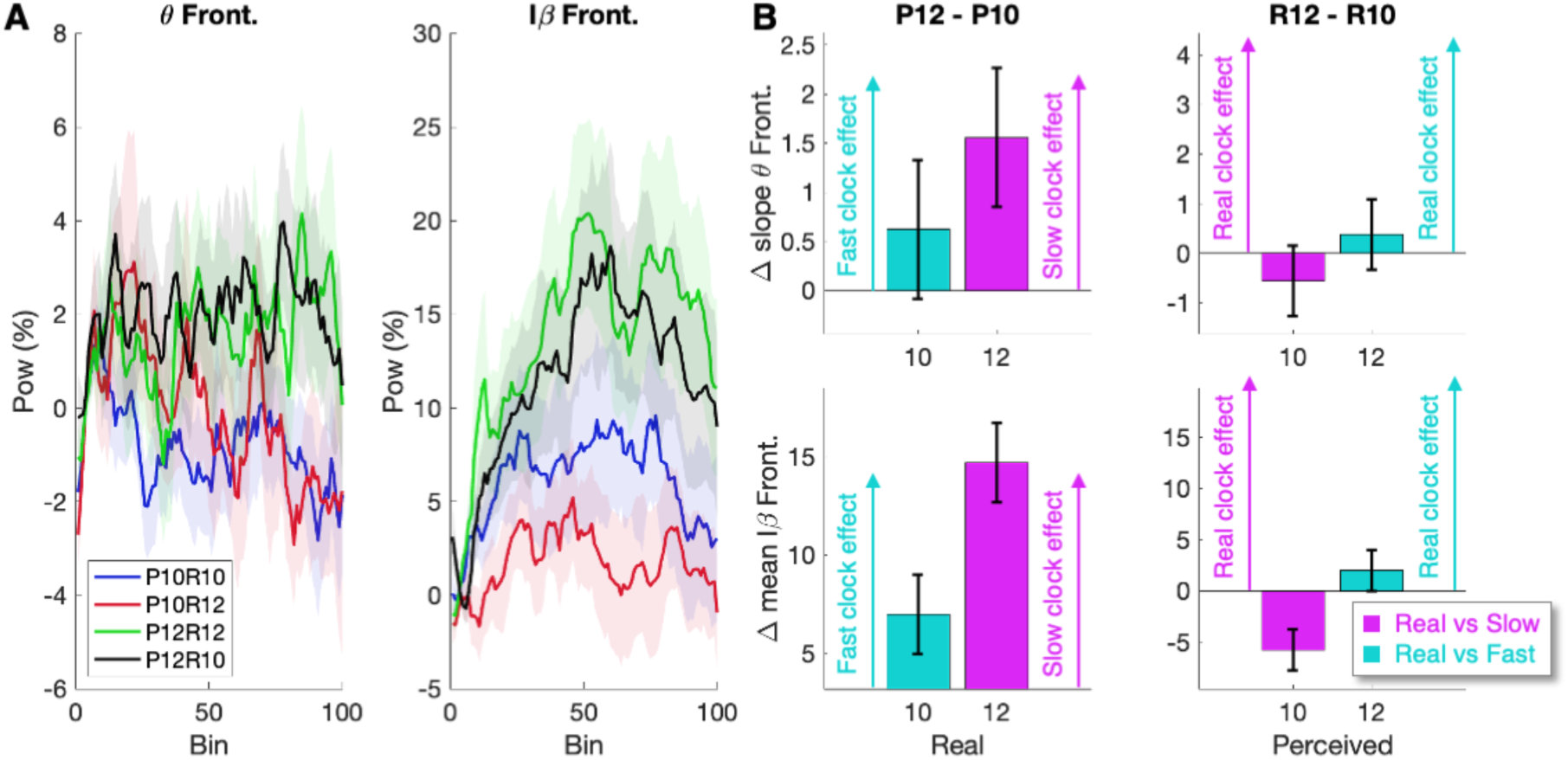
Neural correlates of time deception. A. EEG Power modulations in frontal electrodes. Relative power changes in θ (panel 1) and Iβ (panel 2) over contractions (relative to the first contractions). For both panels, the line represents the averaged distribution, and the shaded areas are the SEM. B. Power differences in the frontal electrodes. Power differences were computed for each contrast. The left panels illustrate perceived-time effects by holding real duration constant and contrasting long versus short perceived durations, whereas the right panels illustrate real-time effects by holding perceived duration constant and contrasting long versus short real durations. The upper panels represent the difference in the slope of the frontal θ power, while the lower panels present the difference in the means of Iβ power over the bins. The bar plots represent model-based estimates of condition differences, with error bars indicating 95% confidence intervals. Intervals not crossing zero indicate statistically significant effects at a Bonferroni-corrected alpha of 0.05.

For Iβ frontal activity, model comparison showed that a second-degree spline model provided a significantly better fit than the linear model (χ²_(4)_ = 207.24, p < .0001) and was therefore retained. In the selected model, a significant main effect of contraction number was observed (F_(2, 45.1)_ = 92.07, p < .0001). No significant main effects of perceived time (F_(1, 9542)_ = 0.41, p = .523) or real time (F_(1, 9542)_ = 0.00, p = .970) were found. A significant interaction between contraction number and perceived time was observed (F_(2, 9542)_ = 39.53, p < .0001), while the interaction between perceived and real time showed a trend (F_(1, 9542)_ = 3.76, p = .052). All other interactions were non-significant (all p > .18). Although the interaction did not reach conventional significance, post-hoc contrasts on the estimated marginal means were conducted to test our a priori hypothesis regarding the influence of perceived time (Schad et al., 2020). They revealed a robust effect of perceived time: Iβ was greater for longer perceived time, both when real time was short (p < .0001) and when it was long (p < .0001), in both cases with very strong effect sizes (respectively, d_z_ = 1.77 and d_z_ = 3.73). Real-time influence on Iβ at short perceived time was significant (p < .0001) and large (d_z_ = 1.46), though Iβ was greater in the short real duration compared to the long real duration. In other words, the difference is contrary to the expected pattern if real duration were to play a role. Finally, there was no effect of real time at long perceived time (p = .051, d_z_ = 0.51, BF = 0.26) (Fig. 3A, 3B). Together, these results suggest that frontal beta dynamic is strongly shaped by perceived time, both when the clock is slowed down and accelerated.

### EEG-EMG Connectivity

After independently investigating the EMG and EEG power modulations, we examined their relationship by computing subject-by-subject correlations (see Methods for details). We first found consistently across participants a negative correlation between EMG and Iβ Cz for all sessions (p < .045, d_z_ > 0.43) (Fig. 4A), indicating more EMG activity for larger Iβ desynchronization. We then investigated the link between EMG and frontal oscillatory activity. Similarly, we found a negative correlation between EMG activity and frontal θ-band oscillations in P10R12 (t_(23)_ = 2.70, p = .013, d_z_ = 0.46), and as a trend in P10R10 (t_(23)_ = 1.99, p = .059, d_z_ = 0.41, BF = 1.14), but no correlations in the P12R12 and P12R10 sessions (p > 0.36, d_z_ < 0.19, BF < 0.32) (Fig. 4B). Conversely, we found a positive correlation between the EMG activity and the Iβ in the P12R10 session (t_(23)_ = 2.25, p = .035, d_z_ = 0.46) and, as a trend, in the P12R12 one (t_(23)_ = 1.75, p = .094, d_z_ = 0.36, BF = 0.80), while there was no correlation in the sessions whose perceived time was shorter (P10, p > 0.75, d_z_ < 0.66, BF < 0.23, for both) (Fig. 4C). These results mirror the trends observed in frontal power modulations (Fig. 3A, 3B), showing lower Frontal-EMG correlations for short perceived times and larger correlations for longer ones, suggesting their dependency on perceived rather than actual time.

**Figure 4.**
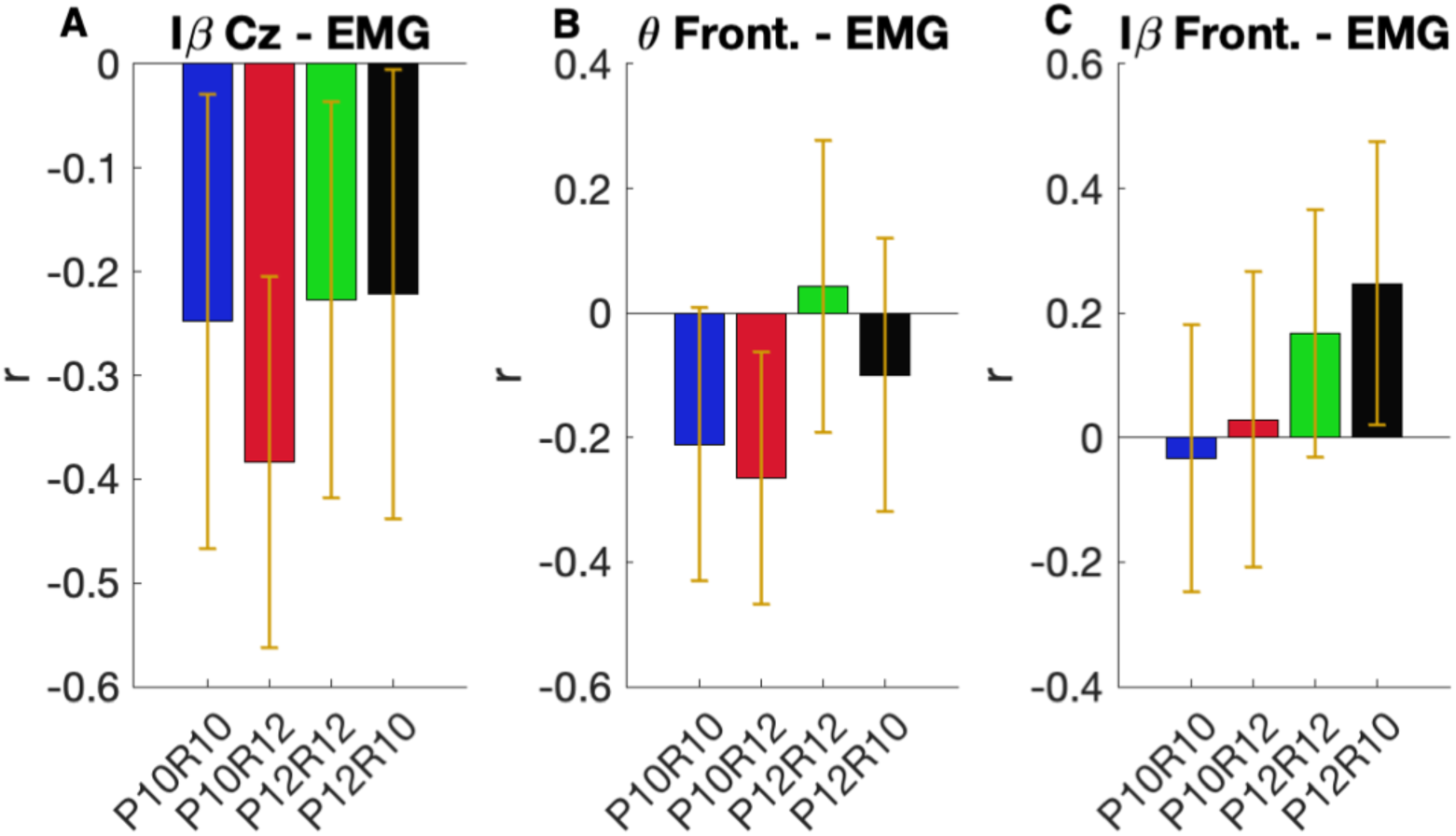
Brain-muscle correlations. Within-subject correlations were performed between EMG and θ Frontal (A), Iβ Cz (B), and Iβ Frontal (C), respectively. The resulting r values were then compared against 0 using t-tests. The bar plots show the average distribution, and the error bars represent the 95% confidence intervals. Intervals that do not cross zero indicate statistically significant effects with p < .05.

### Relationship between frontal oscillatory dynamics and fatigue accumulation

Our results show that perceived time can significantly influence the accumulation of physical fatigue and that the frontal oscillatory dynamics are robustly modulated by perceived rather than actual time. Starting from the assumption that PFC is involved in belief representation (Geuter et al., 2017), we hypothesized that participants with the strongest deceptive effect in the frontal electrodes would also exhibit the largest EMG effect. To test our hypothesis, we split the participants into two groups via a median split computed on the normalized θ-power difference between the two sessions with the same perceived time. The first split separated participants based on the normalized θ-power difference computed between sessions that shared a perceived short duration (P10) but differed in real duration. In a second median split, the same variable was computed for sessions that shared a perceived long duration (P12) but differed in real duration. These power differences were normalized by the difference between the P12R12 and P10R10 conditions to account for the power difference observed under normal clock conditions. In both cases, smaller differences correspond to a larger deceptive effect, as the θ power modulation in frontal electrodes follows the perceived rather than the actual time (‘Subjective-time’ (ST) group). Conversely, participants with larger power differences would rely more on actual than perceived time (‘Objective-time’ (OT) group). Note that we used the θ power difference to perform our median split because of its well-established relationship with cognitive control (Cavanagh & Frank, 2014).

For all subgroups, a linear model with a simple contraction effect (linear) was sufficient, as adding a second-degree spline did not improve model fit (all p > .15). In the ST subgroup defined on the θ P10 difference, significant main effects of contraction number (F_(1, 11)_ = 15.91, p = .0021) and perceived time (F_(1, 4770)_ = 17.03, p < .0001) were observed. Significant interactions were found between contraction number and perceived time (F_(1, 4770)_ = 50.40, p < .0001), between contraction number and real time (F_(1, 4770)_ = 7.05, p = .0079), and a three-way interaction among contraction number, perceived time, and real time (F_(1, 4770)_ = 8.77, p = .0031). Post-hoc contrasts on the estimated marginal slopes indicated that EMG increased more for long perceived time, both when real time was short (p = .014, d_z_ = 0.84) and long (p < .001, d_z_ = 2.05), indicating that EMG can be both slowed down and accelerated, according to the clock calibration. Accordingly, there was no significant influence of real duration when perceived time was short (p = 1, d_z_ = 0.06, BF = 0.29). Finally, when perceived time was long, EMG rate was still modulated by real time (p = <.0001, d_z_ = 1.15) (Fig. 5A, 5B).

**Figure 5.**
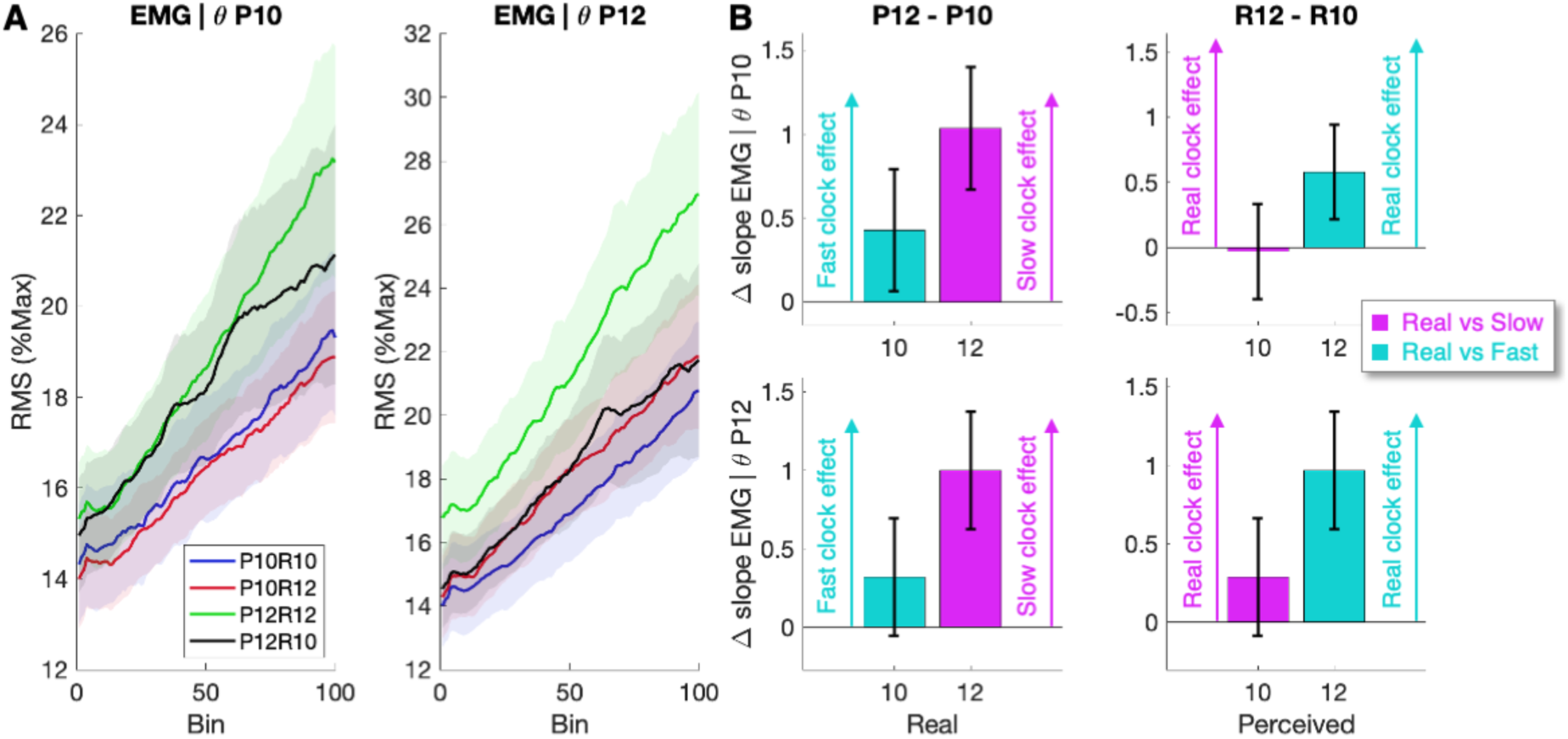
Physical fatigue accumulation in the ‘Subjective-time’ subgroup. A. EMG modulations over the main task. EMG is expressed in normalized RMS over contractions, whose increase reflects neuromuscular fatigue accumulation. Panels contain the ‘Subjective-time’ participants only (N = 12) (based on the deception effects measured in the θ-band, see text for details). Panel on the left was computed on frontal θ absolute sum power difference between sessions sharing the same perceived short duration (P10) and panel on the right between sessions sharing the same perceived long duration (P12). For both panels, the lines represent the averaged distribution, and the error bars are the standard error of the mean. B. EMG differences. EMG differences were computed for each contrast. The left panel illustrates perceived-time effects by holding real duration constant and contrasting long versus short perceived durations, whereas the right panel illustrates real-time effects by holding perceived duration constant and contrasting long versus short real durations. All panels represent differences in the slope of the EMG-RMS across bins. For all panels, the bar plots represent model-based estimates of condition differences, with error bars indicating 95% confidence intervals. Intervals not crossing zero indicate statistically significant effects at a Bonferroni-corrected alpha of 0.05.

In the ST subgroup defined on the θ P12 difference, contraction number (F_(1, 11)_ = 25.13, p < .0001), perceived time (F_(1, 4770)_ = 32.40, p < .0001), and real time (F_(1, 4770)_ = 23.38, p < .0001) showed significant main effects, while interactions between contraction number and perceived time (F_(1, 4770)_ = 38.89, p < .0001), between contraction number and real time (F_(1, 4770)_ = 35.14, p < .0001), and the three-way interaction (F_(1, 4770)_ = 10.39, p = .0013) were also significant. EMG increased more at long perceived time when real time was long (p < .0001, d_z_ = 1.93), supporting a modulation by perceived time under slowed clocks, while the evidence was inconclusive at short real time (p = .13, d_z_ = 0.61, BF = 0.32). Finally, real duration provided inconclusive evidence when perceived time was short (p = 0.22, d_z_ = 0.55, BF = 0.46) and had a significant influence under long perceived time (p < .0001, d_z_ = 1.87) (Fig. 5A, 5B).

In the OT subgroup defined on the θ P10 difference, significant effects of contraction number (F_(1, 11)_ = 33.09, p < .0001), real time (F_(1, 4770)_ = 5.97, p = .015), and an interaction between contraction number and real time (F_(1, 4770)_ = 30.48, p < .0001) were observed, highlighting a dependency on real time only.

Finally, in the OT subgroup defined on the θ P12 difference, main effects of real time (F_(1, 4770)_ = 21.91, p < .0001) and a significant interaction between perceived and real time (F_(1, 4770)_ = 9.91, p = .0017) were observed, along with a significant interaction between contraction number and real time (F_(1, 4770)_ = 4.77, p = .029). Post-hoc contrasts on the estimated marginal means showed that EMG was greater with short perceived time at long real time (p < .0001, d_z_ = 1.31) – that is, in the opposite direction if perceived duration were to play a role, while there was no difference at short real time (p = 0.25, d_z_ = 0.54, BF = 0.29). Similarly, EMG was greater with short real duration at long perceived time (p < .0001, d_z_ = 2.06), while there was no difference at short perceived time (p = 1, d_z_ = 0.21, BF = 0.29).

While this analysis remains exploratory and is partially limited by a median split and a small sample size (n = 12 per group), these results suggest that the frontal power changes play a key role in the effectiveness of the deceptive effect on fatigue accumulation, as measured by EMG. Interestingly, it reveals that, in the ST group, fatigue can be slowed down and accelerated depending on the clock calibration.

### Link between interoceptive awareness and deception

To further explore the individual differences that may affect the strength of the deceptive effect, we considered participants’ interoceptive awareness, measured via the Multidimensional Assessment of Interoceptive Awareness (MAIA) (Mehling et al., 2012) after their last session, as a potential moderating factor. We specifically focused on the ‘Noticing’ and ‘Body Listening’ subscales, which are more closely linked to bodily processes (Iodice et al., 2019). We then tested the correlation between the Noticing and Body Listening scores with the normalized absolute difference of neuromuscular fatigue accumulation (i.e., EMG) for the two sessions with the same subjective time (P10 or P12) but a different real time. In short, a low normalized index indicates no difference between the two sessions, suggesting a large deceptive effect, while a high index reflects the opposite. We hypothesized a positive correlation between the interoceptive score and the normalized index, i.e., the lower the interoceptive awareness, the higher the deceptive effect, and vice-versa. Using non-parametric Kendall’s tau correlation, we found a positive correlation between the Body Listening score and the normalized difference in sessions with the same short perceived time only (τ = 0.29, p = 0.029, unilateral test, H1: positive correlation). No significant correlation was found with the Noticing score (p > 0.27, BF < 0.47). These results suggest that the participant’s body awareness might predict the deceptive effect on neuromuscular fatigue accumulation in the slow-down clock configuration. As our previous findings suggested a role for frontal oscillatory dynamics in the deceptive effect, we tested the correlation between the Noticing and Body Listening scores and the normalized absolute difference in θ frontal power modulations across the two sessions with the same perceived time. We found a positive correlation between the Body Listening score and the θ normalized difference from sessions with the same long perceived time only (τ = 0.31, p = 0.020, unilateral test, H1: positive correlation). While this result supports a link between interoception awareness and the deceptive effect, it remains unclear why this effect appears only when time is accelerated, but not when it is slowed down, as no other significant correlations were found (p > 0.11, BF < 1.02, unilateral test, H1: positive correlation).

## Discussion

In the current study, we used a novel design to test the influence of time deception on physical fatigue accumulation while monitoring associated EEG power modulations. We found that physical fatigue accumulation was modulated by perceived time, specifically accumulating more slowly when the perceived duration was short, independent of real time. Importantly, this deceptive effect was observed in the absence of motivational confounds, demonstrating, for the first time, the direct influence of time deception on the course of physical fatigue. Regarding the neural correlates of the deceptive effect, we found that frontal oscillatory dynamics was largely shaped by perceived time. We then showed that frontal dynamics predict the effectiveness of time deception on fatigue accumulation, suggesting its key role in the time deception effect.

### Physical fatigue follows the perceived time

Our findings reveal that physical fatigue, as indexed by EMG activity increasing across contractions, follows the subjective rather than the objective time when the clock is slowed down, echoing previous work emphasizing the mind’s influence on bodily processes (Craik & Sarbin, 1963; Crum et al., 2011; Crum & Langer, 2007; Langer et al., 2010; London & Monello, 1974; Pomares et al., 2011; Schachter & Gross, 1968; Snyder et al., 1974; Thönes et al., 2021), such as blood glucose level (Park et al., 2016) and healing (Aungle & Langer, 2023). While physical fatigue has been proposed to depend on subjective time (Matta et al., 2024), the current study is the first to accurately disentangle the motivational effects from the temporal deception. Furthermore, our results show an asymmetrical effect of clock manipulation, which might reflect mechanisms of efficiency and homeostatic regulation. Indeed, slowing perceived time may promote more economical effort allocation, delaying fatigue accumulation, and facilitating task completion (Meyniel et al., 2013). In contrast, accelerating perceived time does not produce a corresponding benefit, likely because fatigue functions as a protective signal that limits performance to prevent physiological harm and maintain homeostasis (Noakes, 2012; St Clair Gibson et al., 2003, 2013). As a result, even when perceived time is deceptively accelerated, compensatory mechanisms may be engaged to mitigate fatigue and preserve performance. Yet, complementary analyses (Fig. 5) nuance this effect, showing that some of the participants rely more on perceived time, both when the clock was slowed down and accelerated, while others follow objective time. Our findings suggest that interoceptive awareness may modulate this effect: lower interoceptive awareness is associated with a larger deceptive effect, and vice versa. However, other factors likely influence this perceived-time dependency, such as the strength of participants’ beliefs (e.g., expectations that shorter times lead to less fatigue) (Shiv & Carmon, 2005), the effectiveness of external clock manipulations on the perception of time (Craik & Sarbin, 1963; Thönes et al., 2018), but also participants’ backgrounds or personality (Zhou et al., 2019). Further research is needed to better characterize the factors driving time deception’s effectiveness on fatigue.

### Neural correlates of the time deception effect

In the current study, we found no differences between the normal sessions in the Cz oscillatory dynamics, reflecting activity in motor areas, likely due to a relatively small workload difference (Matta et al., 2025), making any potential difference due to the deceptive manipulation even more challenging to observe. Yet, these parameters were carefully chosen to ensure the implicit plausibility of the biased clock calibration, as increasing the manipulation’s magnitude could have raised awareness of the effect, potentially negating and confounding its impact (Ducrocq et al., 2017; Williams et al., 2014).

Even though the task constraints imposed a relatively short temporal difference between conditions, frontal oscillatory dynamics were significantly shaped by perceived time. Notably, theta-band activity was modulated by perceived time in the slowed-clock condition, whereas beta-band activity was sensitive to perceived time across both slowed and accelerated clocks. Interestingly, both theta and beta-band activity increases have been previously associated with prolonged engagement in demanding tasks (Boksem et al., 2005; Stoll et al., 2016). Specifically, enhanced theta power has been extensively linked to increasing mental (Baumeister et al., 2012; Borghini et al., 2014; Craig et al., 2012; Fan et al., 2015; Käthner et al., 2014; Liu et al., 2010; Shigihara et al., 2013; Tanaka et al., 2014; Wascher et al., 2014) and, albeit less consistently, physical (Bailey et al., 2008; Hottenrott & Gronwald, 2013; Mechau et al., 1998) fatigue – both supposedly sharing common neural resources (Bray et al., 2012; Jin et al., 2024; Marcora et al., 2009; Mehta & Parasuraman, 2014; Van Cutsem et al., 2017). This increase in theta power has been proposed as reflecting a fatigue-related compensatory mechanism (Wascher et al., 2014) to allow sustaining performance, seemingly driven by subjective rather than actual performance. Moreover, given that frontal theta is a well-established marker of cognitive control (Arnau et al., 2024; Cavanagh et al., 2012; Cavanagh & Frank, 2014; Ehrhardt et al., 2022; Howells et al., 2010; Huster et al., 2013; Ishii et al., 1999; Lopez-Gamundi et al., 2024; McFerren et al., 2021; Yu et al., 2022), our results suggest that the cognitive control required to sustain a physical task depends primarily on perceived rather than actual time. Finally, several studies also reported an increase in frontal beta with the rise in fatigue (Bailey et al., 2008; Enders et al., 2016; Hottenrott & Gronwald, 2013; Kubitz & Mott, 1996; Mechau et al., 1998). This increase might also reflect a top-down cognitive control (Buschman & Miller, 2007; Stoll et al., 2016) allowing one to continue the task despite uncomfortable sensations, which might follow perceived rather than actual time. However, our study did not include continuous measurements of subjective experience, such as perceived effort (Pageaux, 2016), which limits our ability to fully characterize these processes. Future studies should incorporate such measures to better understand the relationship between perceived time and participants’ subjective experience of effort. Together, these results highlight the reliance of specific cognitive control time-related correlates to perceived rather than actual time. Additionally, it emphasizes the importance of considering subjective components in cognitive experimental paradigms, as various subjective factors may significantly influence physiological measures.

### A predictive coding perspective

We found that frontal oscillatory dynamics (supposedly reflecting PFC’s activity) and physical fatigue evolve according to perceived rather than actual time. The influence of frontal areas and the PFC on physical fatigue may be explained through predictive coding (Kaptchuk, 2018), a framework suggesting that the brain continuously generates expectations about incoming sensory information (Friston, 2010). When there is a mismatch between expected and actual sensations – a prediction error – the brain can resolve this discrepancy either by updating its expectations (priors) or by actively adjusting bodily states (active inference) (Friston et al., 2009; Pezzulo et al., 2015; Seth, 2013). If we apply this framework to our findings, we may consider beliefs about time as prior factors that regulate physical fatigue through an active inference process. In this situation, the temporal deceptive manipulation induces an inaccurate temporal estimate, which generates a prediction error in the participants about their current fatigue level. In order to minimize such prediction error – the discrepancy between expected and actual fatigue – the brain may modulate interoceptive input (i.e., interoceptive signals related to physical fatigue, later influencing EMG signals), thereby adjusting the evolution of physical fatigue rather than revising its priors (Pagnini et al., 2023). In line with this framework, frontal oscillatory dynamics could reflect temporal priors, as the PFC has been proposed to play a key role in this process (Geuter et al., 2017). Furthermore, previous research has demonstrated that beliefs can influence physiological effector systems such as the autonomic nervous system (Amigo et al., 1993; Meissner, 2009, 2011; Nakamura et al., 2012; Wager et al., 2009). In line with this interpretation, one may speculate that the change in the physical fatigue course is mediated by an indirect pathway that originates in the PFC and regulates peripheral organs (e.g., cardiac, respiratory) (Hassan et al., 2013; Thayer et al., 2009, 2012), impacting muscle fatigue as an outcome to reduce the prediction error. Furthermore, our complementary analysis, splitting the participants based on their PFC power difference between sessions with the same subjective time, support the hypothesis that frontal areas modulate the time manipulation effect. Indeed, as shown in Figure 5, only participants exhibiting strong deceptive effects in the PFC also experienced the deceptive effect on fatigue, as revealed in the EMG recordings. While this observation should be interpreted with caution, given the limited sample size and the exploratory use of a median split, it may pinpoint the PFC as one of the core structures involved in mediating the time deception effects on physical fatigue. Future studies with larger samples and continuous, model-based approaches will be necessary to confirm this relationship and to characterize individual differences in susceptibility to time-deception effects more robustly.

## Conclusion

Our study demonstrates the direct influence of time deception on physical fatigue, irrespective of motivational factors and performance goals, and reveals key neural correlates of this effect. We show that this influence is primarily associated with frontal oscillatory dynamics that closely track perceived time and are pivotal to the effectiveness of such deceptive time manipulation. Building on these findings, we propose potential mechanisms underlying this mind-body interaction. By showing that psychological factors can modulate physical fatigue, our results have broad implications for performance enhancement, but also everyday life and occupational settings, suggesting ways to optimize endurance, productivity, and safety in workers and other physically demanding contexts. Future research should further investigate the neural mechanisms driving this phenomenon from different perspectives (e.g., cardiac, respiratory or fMRI recordings) to further deepen its theoretical understanding. It may also explore applications across various contexts (e.g., sport, work, daily life) and their potential translation into clinical conditions.

## Methods

### Participants

Twenty-eight healthy volunteers with no known history of neurological diseases or physical injuries and regular physical practice (once to several times a week) participated in this study. Out of the 28 individuals, 24 were able to complete the entire protocol and were kept on for the analysis (13 women, age: 24 ± 4, weight: 70 ± 19, mean ± SD). Before participating in the study, all individuals provided informed consent and were informed of their right to withdraw from the study at any time. The four sessions were justified to the participants on the grounds that they allowed (i) the study of fatigue evolution between two different levels of effort (short and long contractions) and (ii) each level to be repeated twice to assess the reproducibility of fatigue measurements at the same level of effort. No further details were provided. At the end of the study, participants completed a survey addressing various aspects of their experience during the sessions (e.g., whether some sessions felt more difficult than others, whether the load seemed consistent across the four sessions, whether recovery time felt the same between sessions, and whether the duration of effort was similar across sessions). No participant reported any specific remarks suggesting awareness of the temporal manipulation. Participants were only informed of the deception after the study was completed. The study procedure was approved by the French Ethical ‘Comité de Protection des Personnes’ (number: 2023-A00412-43).

### Data Acquisition

During each session, all participants were seated on a Biodex isokinetic ergometer with their hips and knees bent to a 90° angle. Their right leg was securely fastened to the ergometer accessory at the ankle with a non-compliant strap just above the malleolus. Two shoulder and one abdominal harnesses guaranteed a reduction in their range of motion and the involvement of other muscles besides the quadriceps. The installation parameters were optimized for each subject and kept identical within subjects across sessions.

Torque and electromyography (EMG) were recorded at 2000 Hz using a Biopac MP150 system and AcqKnowledge software. EMG was recorded through three pairs of electrodes (AG-Cl, diameter: 11 mm, interelectrode distance: 2 cm) placed on the Vastus Medialis, Vastus Lateralis, and Rectus Femoris according to SENIAM recommendations. Before placement, the skin was carefully prepared through shaving and alcohol to ensure low impedance (<5 KΩ). The reference electrode was positioned on the patella of the contralateral knee. EEG was recorded using a 64 electrodes Biosemi Active 2 system with 64 Ag-AgCl sintered active electrodes at 2048 Hz. The EEG electrode locations followed the 10–20 international system. The offset was kept below 20 mV using an active gel. All acquisitions were performed with the light off to minimize electrical noise.

The main task was programmed in Python and displayed through a Windows computer (Resolution: 1920 × 1080; Refresh rate: 60 Hz), using a NiDAQ acquisition card (NI USB-6218, Sample rate: 2000 Hz) to read the torque data (low-pass filtered at 10 Hz) in real-time. The same computer also generated the triggers used to mark the onset and offset of each contraction towards BIOPAC and BIOSEMI, allowing us to synchronize all our signals offline.

### Experimental Protocol

All participants came to the lab on four days (each spaced 3 to 10 days apart) to perform four sessions. On each day, the participants performed the following steps (Fig. 1):

- Warm-up (consisting of 15-20 contractions with increasing intensity, approximately from 20 to 80% of their estimated Maximal Voluntary Torque (MVT))
- 2 Maximal Voluntary Contractions (MVC) (3-4 s each, 30 s spaced)
- 1 MVC, 20 s of rest, 1 Long MVC (30 s), 30 s of rest, 3 minutes of EEG recording at rest (eyes opened)
- 5 minutes of rest
- Main task: 100 contractions (see below)
- 1 MVC (5 s after the 100^th^ contraction of the main task), 20 s of rest, 1 Long MVC (30 s), 30 s of rest, 3 minutes of EEG recording at rest (eyes opened)

Regarding the main task, participants performed 100 contractions at a fixed individualized intensity, defined as 20% of their maximal force, or 40 N.m if their maximal force was larger than 200 N.m. The intensity was the same among the four sessions. While the rest time between contractions was identical for each session (5 s), each contraction’s real (R) and perceived (P) time were independently manipulated. In each session, the contractions’ time was either short (10s) or long (12s), while the clock displayed to the participants was also either short (10 s) or long (12 s). This resulted in a 2 × 2 design with Perceived duration (10 s vs. 12 s) nested within Real duration (10 s vs. 12 s). An accelerated clock was only possible at the shortest real duration, whereas a slowed-down clock was only possible at the longest real duration. This yielded four counterbalanced conditions: P10R10, P12R12, P10R12, and P12R10 (Fig. 1). In practice, a sound indicated to the participants when to begin and end each contraction. Simultaneously with the start sound, a rectangle was displayed in the middle of a black screen to provide online feedback to participants on their actual force relative to the individualized target. The rectangle’s color was updated online based on the ongoing force: red indicated participants were below the targeted force, blue above, and green was in the aimed range (-5% to +5% around the targeted force). A digital clock was displayed in the middle of the colored rectangle 0.75 s after the start of the contraction (preparation time left to the participants to reach the targeted force). This clock allowed us to perform our manipulation, combining real time (R10 or R12) and perceived time (P10 or P12). The clock was therefore either normal (P10R10 and P12R12), slowed down (P10R12), or accelerated (P12R10) without the participants knowing it. After the ending sound, the final perceived time (10.0 or 12.0 s) remained on the screen for 1 s, then the number of contractions already performed was displayed during the resting time. Note that part of the data (only from the two normal sessions, P10R10 and P12R12) has been previously reported (Matta et al., 2025) in the context of a different question; Long MVC were specifically used there.

### Data pre-processing

All data pre-processing and analyses were conducted offline using custom MATLAB (version 2022b) scripts along with EEGLAB functions (Delorme & Makeig, 2004). Torque and EMG data were down-sampled offline to 1000 Hz. Torque data were then low-pass filtered at 10 Hz, while EMG data were band-pass filtered between 20 and 500 Hz and notch filtered between 47 and 53 Hz.

EEG data were down-sampled offline to 250 Hz. A notch filter (47 to 53 Hz) and a high-pass filter with a 1 Hz cut-off frequency were applied. Channels with excessive noise were automatically rejected using the CleanRawData plugin, configured with a 5-second ‘flat line criterion,’ a 0.6 ‘minimum channel correlation,’ and a kurtosis threshold exceeding five times the mean. On average, 61.7 ± 1.8 (mean ± SD) channels were retained for analysis. The electrodes were then re-referenced using a common average reference. Independent Component Analysis (ICA) was performed with the Infomax algorithm, and the resulting components were labeled using the ICLabel plugin (Pion-Tonachini et al., 2019). Components labeled as muscle, eye, heart, line noise, channel noise, or other with a probability above 90% were automatically rejected, averaging 12.9 ± 5.5 components removed. Previously excluded channels were subsequently interpolated.

EEG data from the two resting-state periods were segmented into 2-second non-overlapping windows. For the main task, EEG and EMG data were segmented into 18-second epochs, starting 3 seconds before the contraction onset (as marked by its trigger) and ending 15 seconds after. A final epoch rejection procedure was applied to the main task EEG data to remove additional noise. Epochs with a standard deviation greater than 1.5 times the interquartile range (IQR) above the 3rd quartile or below 1.5 times the IQR under the 1st quartile were rejected for each subject and electrode. This procedure eliminated 3.9% of the epochs over the four sessions.

### Data analysis Force and EMG

#### Pre-Post

The highest torque from the three MVCs performed before the task was used to determine the pre-test Maximal Voluntary Torque (MVT). The post-test measurement was taken as the highest torque recorded during the post-task MVC. The variation rate was calculated as (post – pre)/pre ×100.

#### Main Task

EMG was quantified as the root-mean-square (RMS) during each contraction of the main task from 2 to 10 seconds after the onset of the contraction (Fig. 1B). This period was chosen to exclude the initial contraction phase, when torque stabilization has not yet occurred, and to ensure identical comparison times across all sessions. Within this period, EMG was quantified as the RMS over 180 ms non-overlapping windows, normalized by the RMS EMG associated with the previously selected maximal torque (centered from -90 ms to +90 ms around it) for the respective muscle and session. The three muscle EMG values were averaged over time to yield one EMG value per contraction in the main task.

### EEG Spectral Power

Spectral power was obtained by applying a continuous wavelet transform to the pre-processed, epoched EEG data within the 1 to 50 Hz frequency range. The wavelet length was linearly increased from one cycle at 1 Hz to twenty cycles at 50 Hz, optimizing the tradeoff between temporal and frequency resolution across the entire frequency range. Contractions that were removed were subsequently linearly interpolated.

Spectral power was specifically extracted for the Theta (θ, 4-8 Hz) and an Individualized Beta band (Iβ), which was defined for each subject to account for inter-individual variability in the beta desynchronization-specific frequency band. Specifically, the frequency with the largest desynchronization was identified for each subject, and Iβ was defined as the range from -2 to +2 Hz around this frequency (Matta et al., 2025). Power from the lateral-frontal (F7-F8) and mid-frontal (F3-F4) electrodes was averaged to create a ’Frontal’ cluster, chosen to reflect activity in the ventrolateral and dorsolateral prefrontal cortex (Okamoto et al., 2004; Robertson & Marino, 2015). The Cz electrode was used to reflect motor cortex activity associated with quadriceps contraction, a location confirmed by topographically visualizing beta desynchronization (Dal Maso et al., 2012; Forrester et al., 2006; Gordon et al., 2023; Meier et al., 2008).

#### Pre-Post Analysis

Raw spectral powers from pre- and post-EEG resting state recordings were independently averaged over epochs (90) and time (0 to 2 s). The variation rate was calculated between the pre- and post-values for each frequency band and region of interest.

#### Main Task Analysis

For each contraction, raw spectral power bands during the period of interest (2 to 10 s) were normalized to the first contractions of the session, averaged over the same 2 to 10 s period (Fig. 1B). This normalization, focusing on the fatigue effect, involved subtracting the baseline and dividing by it, then multiplying by 100 to express power values as a percentage of a non-fatigued state. The baseline was defined as the average of contractions 2 to 5, excluding the first contraction due to its significant torque variability in all sessions. The resulting values were averaged over time and frequencies, yielding one power value per frequency band, contraction, and region of interest.

### EEG-EMG Connectivity

To investigate the EMG-EEG relationships, we computed Pearson’s correlation between EMG and Iβ Cz, θ Frontal, and Iβ Frontal, for each subject, respectively. Each variable was independently averaged in bins of 10 contractions. We then performed bilateral one-sample t-tests to determine whether the r values significantly differed from zero.

### Statistical analyses

Dependent variables were analyzed using linear mixed-effects models (LMMs) to account for repeated measurements across participants. For variables measured longitudinally across 100 contractions, contraction number was included as a continuous predictor, z-scored within participants, and modeled either linearly or using a second-degree natural spline to capture potential non-linear trends. Real time and perceived time were included as fixed factors, and participants were modeled with a random intercept and an uncorrelated random slope for contraction number. Model comparisons were performed using likelihood-ratio tests to select the best degree of complexity for the contraction term (i.e., the degree of the natural splines function), yielding models of the form: *DependentVar ∼ ns(NumContraction_z, df = d) * Perceived * Real + (1 + NumContraction_z || Participant)*, where *ns* represents the natural cubic spline function applied to the *NumContraction_z* factor, and d represents its degree, which was set to either 1 or 2 according to the best model fit (note that d = 1 represents a linear model, d > 1 a non-linear model). In the above formulation, the asterisks denote that the fixed terms are considered with all interactions, while we consider the slope of the *NumContration_z* factor for the random terms.

Our primary aim was to assess the rate of change (slopes) across the session. Specifically, we were interested in the effect of perceived time at short and long real durations, and conversely, the effect of real time at short and long perceived durations, yielding to four contrasts of interest. To test these effects, we adopted a hierarchical approach to post-hoc contrasts. When a three-way interaction between contraction number, perceived time, and real time was significant, this indicated that the rate of change depended jointly on both perceived and real time; in this case, post-hoc contrasts were conducted using estimated marginal trends to extract slopes for the four contrasts of interest, directly addressing our primary hypothesis concerning changes in rate. If only the two-way interaction between perceived and real time was significant, indicating a modulation of overall levels rather than slope differences, post-hoc contrasts were conducted on estimated marginal means for the corresponding four contrasts. If neither interaction was significant, post-hoc contrasts were not reported. In all cases, contrasts were adjusted for multiple comparisons using a Bonferroni correction.

For variables measured as pre-post differences, LMMs included perceived and real time as fixed factors, with participants modeled as a random intercept, resulting in models of the form:

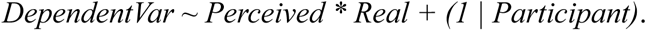

In the case of non-significant results in the post hoc contrasts, Bayes Factors (BFs) were also computed to provide additional evidence regarding the presence or absence of an effect. In the Bayesian framework, the BF represents the ratio of the likelihood of the data under the alternative hypothesis to that under the null hypothesis. Conventionally, BFs > ∼3 provide evidence in favor of the alternative hypothesis, while BFs < ∼0.3 indicate evidence in favor of the null hypothesis (van Doorn et al., 2021).

Statistical analyses were conducted using R (CranR, version 4.3.1).

## Data availability statement

The datasets will be made publicly available on the Open Science Framework (OSF) upon publication of the manuscript.

## Ethical statement

The study procedure was approved by the French Ethical ‘Comité de Protection des Personnes’ (number: 2023-A00412-43).

## Acknowledgements

This project has received funding from the H2020 European Research Council (ERC) under the European Union’s Horizon 2020 research and innovation programme (grant agreements No 101075930 – Andrea Alamia). Views and opinions expressed are however those of the author(s) only and do not necessarily reflect those of the European Union or the European Research Council. Neither the European Union nor the granting authority can be held responsible for them. Pierre-Marie Matta received funding from the French Ministère de l’Enseignement Supérieur et de la Recherche (MESR). We thank Maxime Picquet, Romain Martinie, Khiara Vorzillo, and Nina Bertho for their valuable help with the data acquisition. We are also grateful to Laurent Gonthier for his technical support in setting up the online force acquisition card, and to Nathalie Vayssiere for her assistance in obtaining the ethical agreement.

